# Testosterone induces a conditioned place preference to the nest of a monogamous mouse under field conditions

**DOI:** 10.1101/2021.01.03.425165

**Authors:** Radmila Petric, Matina C. Kalcounis-Rueppell, Catherine A. Marler

## Abstract

Transient increases in testosterone (T-pulses) occur after social interactions in males of various vertebrate species, but the functions of T-pulses are poorly understood. Under laboratory conditions, the rewarding nature of T-pulses induces conditioned place preferences (CPPs), but what are the effects in a complex field environment? We present the first evidence that T-pulses administered to males at their nest site in the wild increased time spent at the nest regardless of pup presence in the monogamous, biparental, and territorial California mouse (*Peromyscus californicus*). Female partners of the T-males, in turn, spent less time at the nest. Independent of treatment, mice produced more ultrasonic vocalizations (USVs) when alone, but T-mice produced more USVs than controls. T-males produced USVs with a smaller bandwidth that likely traveled farther. Our combined results provide compelling evidence that T-pulses can significantly shift the behavioral focus and location of individuals in a complex field setting.

## Introduction

Animals frequently adjust their allocation of time as they move through various life-history stages and meet different social challenges. One mechanism for adjusting approach to a stimulus is through rewarding or reinforcing neural processes (Glickman and Schiff 1967) through the repeated linkage between testosterone (T) release and the presence of a stimulus. T-pulses can act as an internal reward (Gleason et al. 2009) or reinforcing stimulus that when released naturally or through an injection can increase approach to different stimuli such as the physical location in which the T-pulse was experienced, at least under laboratory conditions in rodents (e.g. Zhao and Marler 2014). Because male T-pulses are released in response to social conditions such as a male-male challenge or a male-female interaction across a variety of species including humans (Gleason et al. 2009), we speculate that neural reward and reinforcement mechanisms allow adjustment to changing social challenges that can be linked to location (Zhao and Marler 2016), such as different areas within a territory. In the case of a biparental species, T release near the nest may provide a mechanism for increasing a male’s attendance at the nest, as suggested by the results of a laboratory study (Zhao and Marler 2014). Reinforcing effects can alter the probability for the successful acquisition of essential resources necessary for survival and reproduction (Tinbergen 1957). One paradigm for testing reinforcing effects is to assess changes in behavioral preference for a location at which the stimulus (i.e. T-pulse in response to a social stimulus; Gleason et al. 2009) occurred in the form of a conditioned placed preference (CPP) (Arnedo et al. 2000; Frye et al. 2001). The reinforcing effects occur via activation of the internal reward system (Bell and Sisk 2013). We explore the hormone T as a stimulus that has rewarding/reinforcing effects (Arnedo et al. 2000; Frye et al. 2001; Zhao and Marler 2014; 2016; Zhao et al. 2019; 2020), albeit a weak effect compared to drugs of abuse (Roozen et al. 2004).

The “Challenge Hypothesis” states that male-male encounters induce increases in T in response to challenges from other males (Wingfield et al. 1990). In California mice (*Peromyscus californicus*), T-pulse release occurs after male-male aggressive encounters which influence future behavior under laboratory conditions (Zhao and Marler 2014; 2016). Males that previously won a dispute can form CPPs for the encounter location (Gleason et al. 2009). In monogamous species, the formation of CPPs can be dependent on the familiarity of the environment and the pair-bond status (Zhao and Marler 2014; 2016). For example, in pair-bonded California mice, T-pulses induce CPPs in familiar but not unfamiliar environments (Zhao and Marler 2014; 2016). Interestingly, the opposite was true for sexually naïve males, in which T-pulses induced CPPs in unfamiliar but not in familiar environments (Zhao and Marler 2014; 2016). Therefore, these T-pulses can influence both social interactions and location preferences.

T-pulses modulate other behaviors such as vocalizations (Pultorak et al. 2015; Remage-Healey and Bass 2006), which can affect aspects of sexual selection. Within minutes of a T-pulse in Gulf toadfish (*Opsanus beta*) and plainfin midshipman fish (*Porichthys notatus*), males increase call rate and duration of calls which females tend to prefer (Remage-Healey and Bass 2004; 2006). Male California mice administered a single T-pulse and placed in the presence of a novel female decrease production of calls associated with courtship in pair-bonded but not unpaired mouse males in the laboratory (Pultorak et al. 2015). This finding indicates that in this species, T-pulses reduce extra-pair mating effort by inhibiting the production of courtship calls to unfamiliar females and acting as a fidelity mechanism (Pultorak et al. 2015). T-pulses also have long-term effects on call production in California mice, such that days after multiple T-pulse administration in the field, males produced more call types with a trend to produce more ultrasonic vocalizations (USVs) (Timonin et al. 2018).

While single and multiple T-pulses can alter behavior, the effect of T-pulses on location preferences have only been examined in controlled and simplified laboratory conditions with few competing behavioral choices available, as would occur in the field (Hurley and Kalcounis-Rueppell 2018). There is also a knowledge gap regarding long-term effects (greater than 24 hours) of multiple T-pulses, especially how T-pulses alter time allocation and vocal behavior in complex field settings characterized by multiple and competing biotic and abiotic cues. We hypothesized that, in the field, T-pulses would reinforce behaviors in the area where the social experiences induced T-pulses through the formation of CPPs that would, in turn, alter associated social behavior. Here we tested three predictions: 1) pair-bonded males receiving T-injections at the nest, would spend more time at the nest; 2) females would adjust for the increased time that her T-injected mate spent at the nest by decreasing her time at the nest and allocating more time to activities away from the nest (based on Trainor and Marler 2001); 3) T-pulses would induce changes in type and number of USVs produced as part of both the direct effects of T on behavior and the indirect effects on the pairs’ social adjustment to the altered time allocation to a specific location (Timonin et al. 2018).

We tested our hypothesis in the well-studied monogamous and territorial California mouse, by administering three T-pulses to paired males at the nest site (Figure 1; see methods for details). In this species, males balance their time between mate attendance, offspring care, and territory defense (Gubernick and Alberts 1987; Gubernick et al. 1993; Gubernick and Teferi 2000), and the various behaviors influenced by T. In the laboratory and the wild, California mouse adults frequently produce USVs. In the field, sustained vocalizations (SVs) and barks are frequently recorded (Briggs and Kalcounis-Rueppell 2010; Kalcounis-Rueppell et al. 2018; Timonin et al. 2018), but in the laboratory, additional USVs have been recorded including simple and complex sweeps (Pultorak et al. 2015; 2017; 2018; Rieger and Marler 2018; see USVs details in the methods). SVs are the most common call type recorded in the field as single calls or bouts of multiple calls that are categorized based on the number of calls in a bout (1SV, 2SV, 3SV, 4SV) and are long, low modulation SVs with harmonics (Kalcounis-Rueppell et al. 2018). The SVs often occur when a member of a pair is greater than 1m away and under these conditions may serve as long-distance contact vocalizations, possibly to maintain the pair-bond (Briggs and Kalcounis-Rueppell 2011). Free-living California mice maintain strict territories (Ribble and Salvioni 1990), therefore, social interactions at the nest occur primarily between pair members and include production of SVs. This is consistent with the common production of SVs between pairs in the laboratory (Pultorak et al. 2018). Thus, the monogamous reproductive system of the California mouse and their known time management and vocalization behaviors contribute to a compelling system for assessing behavioral responses to T-pulses and the establishment of male T-induced CPP in the field to alter the amount of time that males spend at the nest.

**Figure 1.**
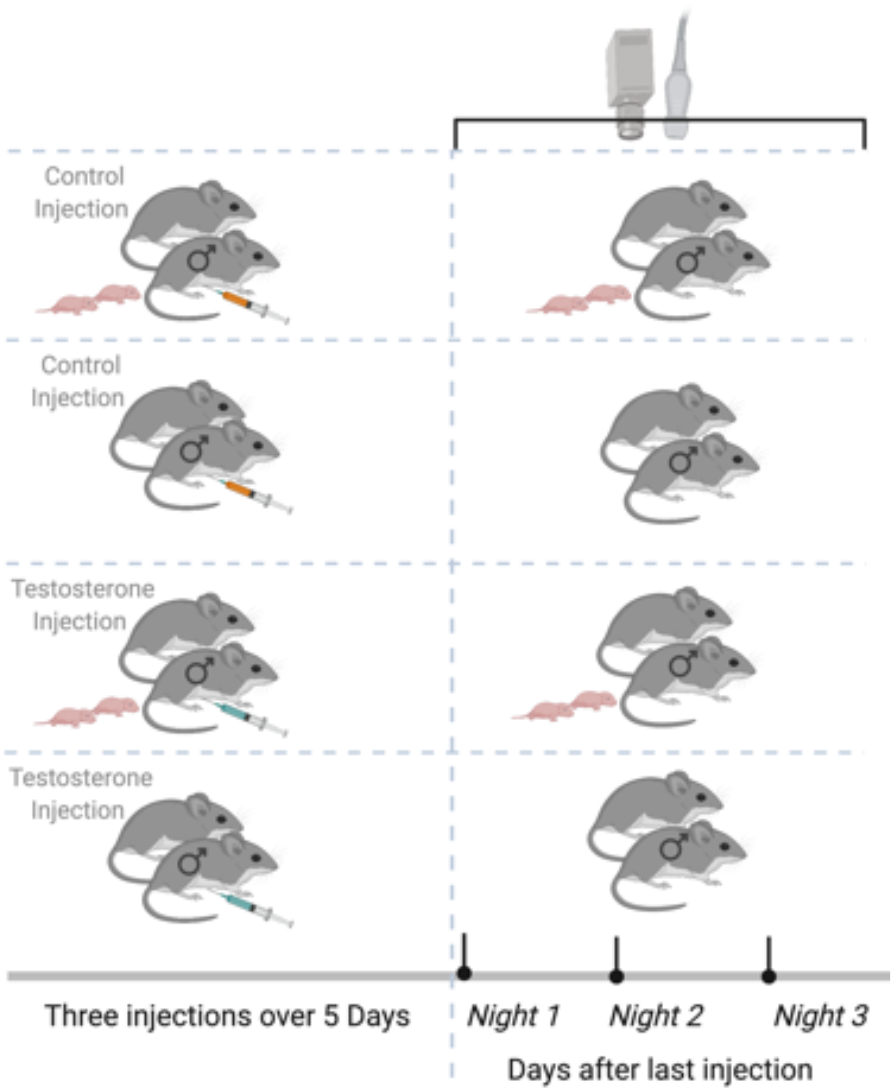
Experimental Design. Paired California mouse males with and without pups were randomly assigned to receive three subcutaneous injections over five nights. After the third and last injection, we deployed the remote sensing equipment (automated radio telemetry, audio recording, and thermal imaging) to record individual behaviors for three consecutive nights. Created with biorender.com

## Results

### Time at the

Overall, T-males spent 14% more time at the nest than C-males (GLMM Estimate 0.14±0.05, p=0.02; Figure 2A). The T-conditioning appeared to be additive in the response to pups where T-males with pups spent 23%o more time at the nest than T-males without pups (GLMM Estimate 0.21±0.04, p<0.01; Figure 2B; Supplemental Table 1).

**Figure 2.**
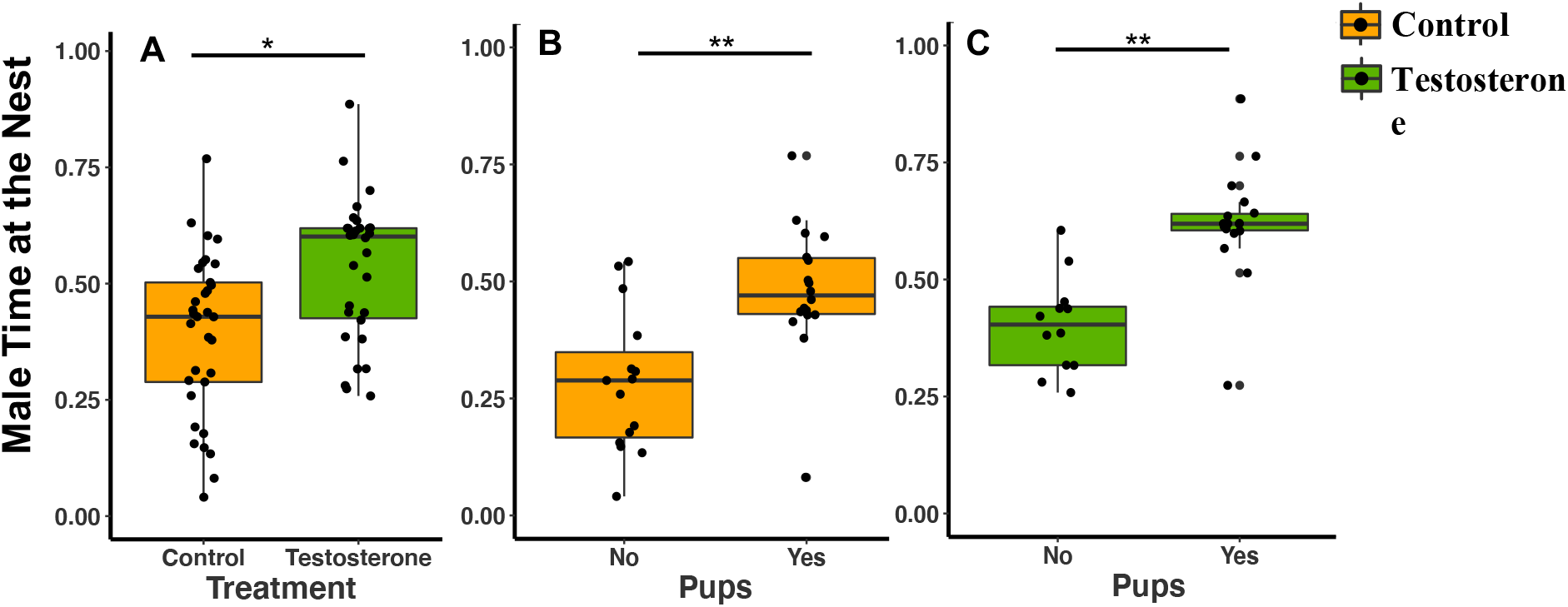
California Mouse (*Peromyscus californicus*) Male Time at the Nest by Treatment Type. **A)** Male time at the nest by treatment type (T=10 and C=11). T-males spent 14% more than C-males (GLMM Estimate 0.14±0.05, p=0.02). **B)** Control male time at the nest by pups (C with pups, n=6; C without pups, n=5). **C)** Testosterone male time at the nest by pups (T with pups, n=6; T without pups). T-males with pups spent 15% more time at the nest than C-male with pups, and T-males without pups spent 12% more time at the nest than C-males without pups (treatment GLMM Estimate 0.13±0.03, p<0.01; pups GLMM Estimate 0.21±0.03, p<0.01). Circles represent individual data points.

**Supplemental Table 1.** Descriptive Statistics on California Mouse Male Time at the Nest, Presence of Pups at the Nest, Season, Body Mass, Number of Nights Required to Administer Three Inj ections, and Recording Night After the Last Injection by Treatment Type. Each male received three testosterone (T=10) or saline (C=11) injection at the nest, after the final inj ect, we recorded time spent at the nest for three consecutive nights.

|  |  | **n** | **Mean** | **SE** | **Median** | **Min** | **Max** |
| --- | --- | --- | --- | --- | --- | --- | --- |
| **Time at the Nest** | C | 33 | 0.390 | 0.030 | 0.430 | 0.040 | 0.770 |
|  | T | 30 | 0.530 | 0.030 | 0.600 | 0.260 | 0.890 |
| **Time at the Nest and Pups** | C with Pups | 18 | 0.480 | 0.030 | 0.470 | 0.080 | 0.770 |
|  | T with Pups | 18 | 0.630 | 0.030 | 0.620 | 0.270 | 0.890 |
|  | C without Pups | 15 | 0.280 | 0.040 | 0.290 | 0.040 | 0.540 |
|  | T without Pups | 12 | 0.400 | 0.030 | 0.400 | 0.260 | 0.610 |
| **Time at the Nest and Season** | C Season One | 21 | 0.344 | 0.040 | 0.313 | 0.041 | 0.769 |
|  | T Season One | 12 | 0.549 | 0.054 | 0.587 | 0.258 | 0.886 |
|  | C Season Two | 12 | 0.476 | 0.034 | 0.491 | 0.178 | 0.631 |
|  | T Season Two | 18 | 0.523 | 0.031 | 0.601 | 0.281 | 0.700 |
| **Body Mass** | C | 33 | 39.050 | 0.707 | 40.000 | 30.500 | 43.000 |
|  | T | 30 | 38.700 | 0.370 | 38.750 | 36.000 | 42.000 |
| **Injection Night** | C | 11 | 3.273 | 0.141 | 3.000 | 3.000 | 4.000 |
|  | T | 10 | 3.100 | 0.100 | 3.000 | 3.000 | 4.000 |
| **Time at the Nest and Recording Night** | C - Night One | 11 | 0.366 | 0.058 | 0.414 | 0.134 | 0.631 |
|  | T - Night One | 10 | 0.490 | 0.052 | 0.569 | 0.258 | 0.666 |
|  | C - Night Two | 11 | 0.375 | 0.045 | 0.429 | 0.081 | 0.552 |
|  | T - Night Two | 10 | 0.555 | 0.044 | 0.604 | 0.317 | 0.763 |
|  | C - Night Three | 11 | 0.435 | 0.056 | 0.435 | 0.041 | 0.769 |
|  | T - Night Three | 10 | 0.554 | 0.053 | 0.564 | 0.317 | 0.886 |

Male time at the nest was not statistically influenced by season (GLMM Estimate −0.09±0.06, p=0.17), body mass (GLMM Estimate −0.01±0.01, p=0.51), total nights needed to administer all three injections (GLMM Estimate −0.09±0.08, p=0.26) or recording night (night two GLMM Estimate 0.04±0.04, p=0.39; night three GLMM Estimate 0.07±0.04, p=0.11; nights two and three are compared to night one after final injection for this analysis).

Females were not subjected to T-injections, but we examined their responses to their T-injected mates. T-females spent 15% less time at the nest than C-females (GLMM Estimate −0.16±0.06, p=0.02; Figure 3A). Females with pups in the nest spent more time at the nest than females without pups (pups GLMM Estimate 0.19±0.06, p<0.01; Figure 3B and 3C; Supplemental Table 2).

**Figure 3.**
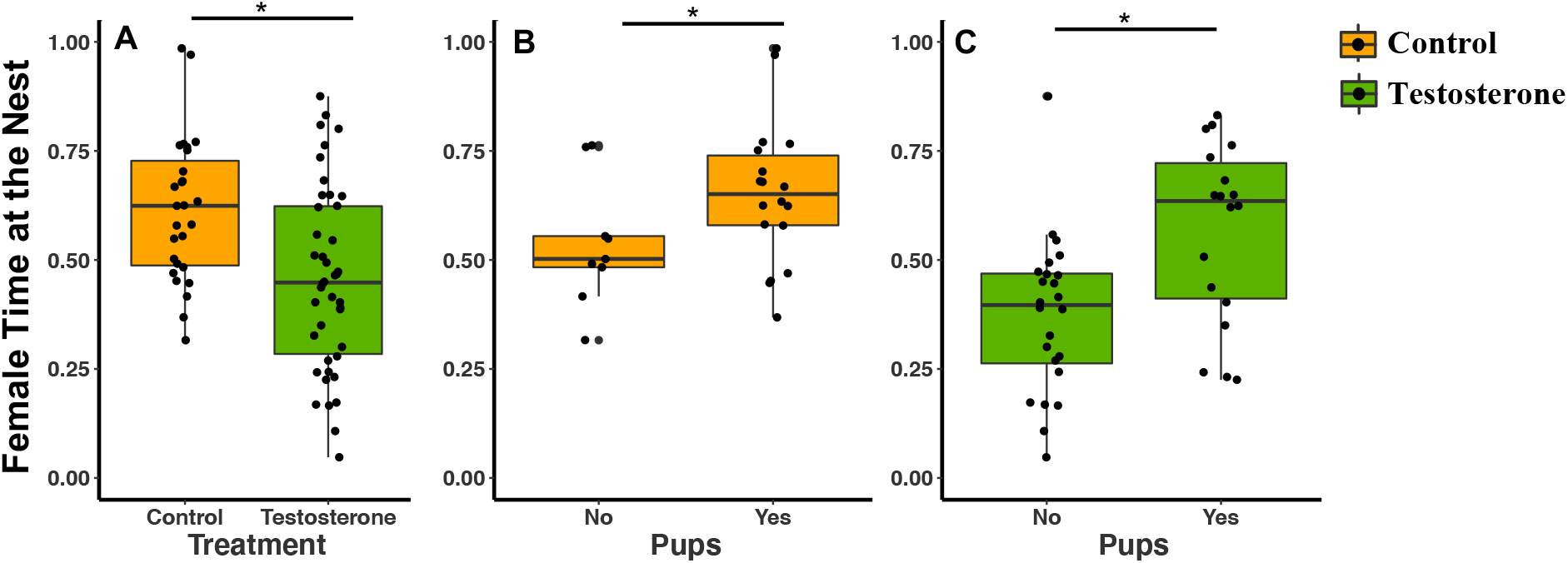
California Mouse Female Time at the Nest by Treatment Type. **A)** Female time at the nest by treatment type (T = 14 and C=9). T-females spent 15.8% less time at the nest than C-females (GLMM Estimate −0.16±0.06, p=0.02). **B)** Control female time at the nest by pups (C with pups, n=6; C without pups, n=3). **C)** Testosterone female time at the nest by pups (T with pups, n=6; T without pups, n=8). When we consider both treatment and pupus, there was a significant effect of pups on female time at the nest (pups GLMM Estimate 0.54±0.24, p<0.04), but there was no treatment effect (treatment GLMM Estimate −0.05±0.25, p=0.84). C-females with pups spent 11.6% more time at the nest than C-female without pups. T-females with pups spent 19.4% more time at the next than T-females without pups. Circles represent individual data points.

**Supplemental Table 2.** Descriptive Statistics on California Mouse Female Time at the Nest, Presence of Pups at the Nest, Season, Body Mass, Number of Nights Required to Administer Three Injections, and Recording Night After the Last Injection by Treatment Type. Each female was paired with a male who received three testosterone (T=14) or saline (C=9) injection at the nest, after the final inject, we recorded time spent at the nest for three consecutive nights.

|  |  | **n** | **Mean** | **SE** | **Median** | **Min** | **Max** |
| --- | --- | --- | --- | --- | --- | --- | --- |
| **Time at the Nest** | C | 27 | 0.614 | 0.032 | 0.624 | 0.316 | 0.985 |
|  | T | 42 | 0.456 | 0.033 | 0.448 | 0.048 | 0.875 |
| **Time at the Nest and Pups** | C with Pups | 18 | 0.653 | 0.039 | 0.651 | 0.369 | 0.985 |
|  | T with Pups | 18 | 0.567 | 0.048 | 0.635 | 0.225 | 0.831 |
|  | C without Pups | 9 | 0.537 | 0.049 | 0.502 | 0.316 | 0.763 |
|  | T without Pups | 24 | 0.373 | 0.037 | 0.397 | 0.048 | 0.875 |
| **Time at the Nest and Season** | C Season One | 12 | 0.557 | 0.034 | 0.526 | 0.417 | 0.771 |
|  | T Season One | 24 | 0.39 | 0.043 | 0.338 | 0.048 | 0.875 |
|  | C Season Two | 15 | 0.66 | 0.048 | 0.679 | 0.316 | 0.985 |
|  | T Season Two | 18 | 0.546 | 0.044 | 0.501 | 0.108 | 0.832 |
| **Body Mass** | C | 27 | 43.06 | 0.85 | 43 | 34.5 | 51 |
|  | T | 42 | 41.93 | 0.752 | 41 | 35 | 51 |
| **Time at the Nest and Recording Night** | C - Night One | 9 | 0.566 | 0.057 | 0.555 | 0.316 | 0.771 |
|  | T - Night One | 14 | 0.409 | 0.065 | 0.382 | 0.048 | 0.832 |
|  | C - Night Two | 9 | 0.649 | 0.054 | 0.625 | 0.452 | 0.971 |
|  | T - Night Two | 14 | 0.425 | 0.051 | 0.42 | 0.166 | 0.875 |
|  | C - Night Three | 9 | 0.628 | 0.057 | 0.624 | 0.417 | 0.985 |
|  | T - Night Three | 14 | 0.536 | 0.051 | 0.509 | 0.225 | 0.801 |

T-females without pups spent 19.4% less time at the nest than T-females with pups, whereas C-females without pups spent 11.6% less time at the nest than C-females with pups (Supplemental Table 2). T- and C-females spent more time at the nest on night three of recording compared to night one of recording (night three GLMM Estimate 0.10±0.04, p<0.02; Supplemental Table 2). T-females spent 13% more time on night three than night one and C-females spent 6% more time on night three than night one (Supplemental Table 2). Females spent less time in the nest during season one than season two, independent of treatment (season one GLMM Estimate −0.15±0.06, p=0.02; Supplemental Table 2). T-females spent 15.6% less time at nest during season one than season two (Supplemental Table 2). While C-male time at the nest was not influenced by season, C-females spent 10.3% less time at the nest during season one than season two (Supplemental Table 2). Female time at the nest was not influenced by body mass (GLMM Estimate 0.01±0.01, p=0.24) or mass difference between the female and the male (GLMM Estimate 0.01±0.01, p=0.17). We also examined within pair comparisons and found a significant negative effect of T and male time at the nest on female time at the nest (T GLMM Estimate −0.15±0.07, p=0.04, Time at the Nest GLMM Estimate 0.36±0.17, p=0.04; Supplemental Table 3). T-females spent 5% less time at the nest than their mates, whereas, C-females spent 18% more time at the nest than their mates (Supplemental Table 3).

**Supplemental Table 3.** Descriptive Statistics on California Mouse Pair Time at the Nest. Each female was paired with a male who received three testosterone (T=10) or saline (C=8) injection at the nest, after the final inject, we recorded time spent at the nest for three consecutive nights.

|  |  | n | Mean | SE | Median | Min | Max |
| --- | --- | --- | --- | --- | --- | --- | --- |
| **Pair Time at the Nest** | C Female | 24 | 0.610 | 0.036 | 0.580 | 0.316 | 0.985 |
|  | T Female | 30 | 0.495 | 0.036 | 0.470 | 0.108 | 0.832 |
|  | C Male | 24 | 0.435 | 0.025 | 0.440 | 0.147 | 0.631 |
|  | T Male | 30 | 0.533 | 0.028 | 0.601 | 0.258 | 0.886 |

### USVs at the Nest

We recorded a total of 549 USVs across the 26 nest sites (T USVs=368, C USVs=181). We assigned context to 385 USVs from video; 157 USVs were produced when a mouse was alone (T USVs=101, C USVs=56), 119 USVs were produced when the mouse was <1m away from another mouse (T USVs=94, C USVs=25), and 109 USVs were produced when the mouse was >1m away from another mouse (T USVs=76, C USVs=33). T-pairs produced twice as many total USVs at the nest than C-pairs (GLMM Estimate 0.87±.40, p=0.04; Figure 4A; Supplemental Table 4). Independent of treatment, pairs also produced more USVs on night one than night three after the last injection, (night two GLMM Estimate −0.33±0.26, p=0.15; night three GLMM Estimate −0.76±0.26, p=0.01; Figure 4B and 4C; Supplemental Table 4). The effect of T on number of USVs could either be a product of direct effects of T on USVs or indirectly because of more time spent apart. Limits on sample size make this difficult to determine (but see Supplemental results Figure 1 and see results for specific call types below).

**Figure 4.**
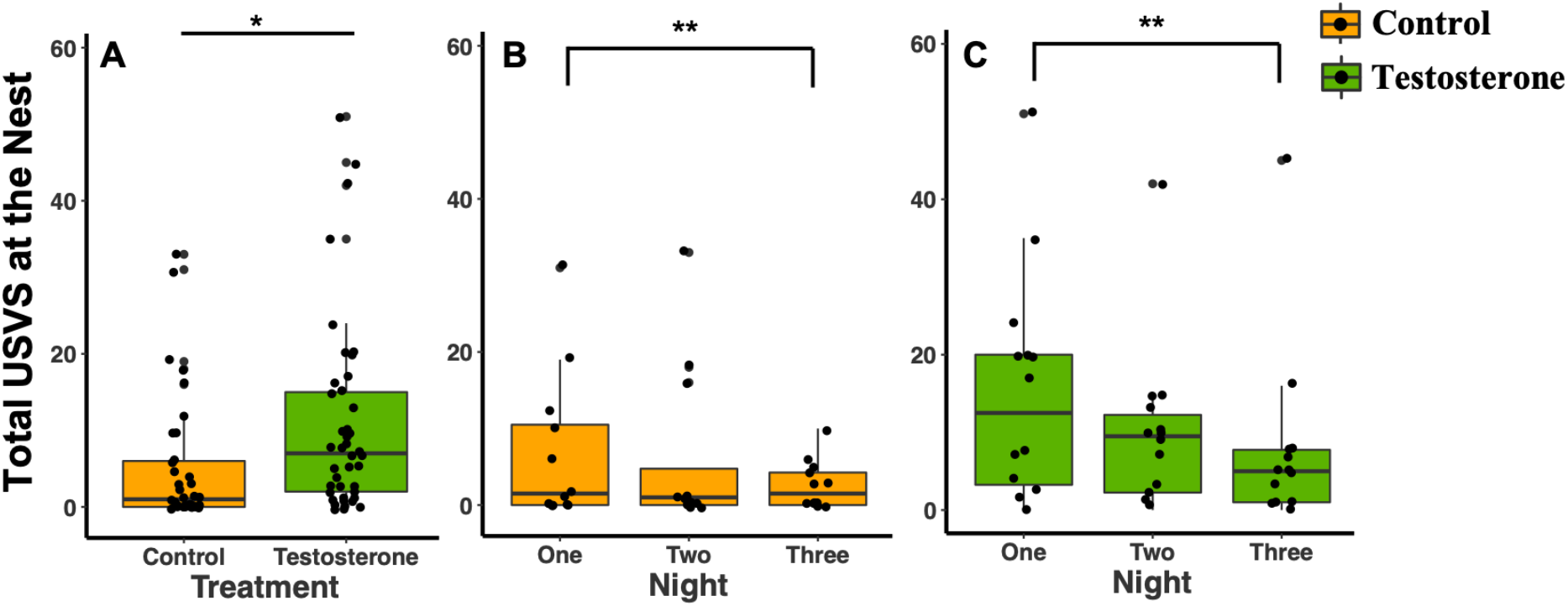
Total USVs Produced at the Nest. A) Total USVs by treatment type. B) C dyad total USVs by night (C=I2). **C)** T dyad total USVs by night (T=I4). Male-female dyads produced more total USVs at T-nests than C-nest (GLMM Estimate 0.87±0.40, p=0.04). Both treatment type and night three post injection significantly influenced total USVs produced at the nest (treatment GLMM Estimate 0.91 ±0.42, p=0.04; night three GLMM Estimate −0.76±0.26, p<0.01). Circles represent individual data points.

**Figure 5.**
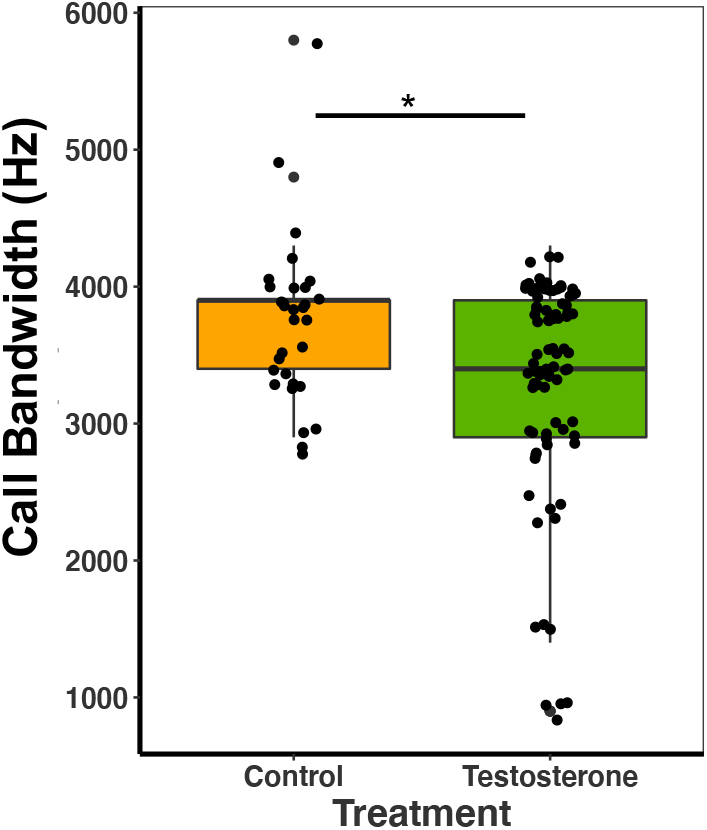
Median and quantiles of call bandwidth (Hz) for male California mouse. The first call in the sequence for 1-, 2-, 3- and 4SVs produced by males (Testosterone = 86 and Control=31). Testosterone males produced calls with a 11.25% smaller bandwidth than control males (GLM Estimate −580.22±182.47, p<0.01). Circles represent individual data points.

**Supplemental Table 4.** Descriptive Statistics on California mouse USVs Produced Per Minute and Per Night at the Nest, Presence of Pups at the Nest, Season, Body Mass, Number of Nights Required to Administer Three Injections, and Recording Night After the Last Injection by Treatment Type. Each male received three testosterone (T=14) or saline (C=12) injection at the nest, after the final inject, we recorded USVs at the nest for three consecutive nights.

| **USVs at the Nest** |  | **n** | **Mean** | **SE** | **Median** | **Min** | **Max** |
| --- | --- | --- | --- | --- | --- | --- | --- |
| **USVs** | C | 36 | 5.083 | 1.421 | 1.000 | 0.000 | 33.000 |
|  | T | 42 | 10.810 | 1.942 | 7.000 | 0.000 | 51.000 |
| **USVs and Pups** | C with Pups | 21 | 3.381 | 1.096 | 1.000 | 0.000 | 18.000 |
|  | T with Pups | 15 | 10.600 | 4.154 | 3.000 | 0.000 | 51.000 |
|  | C without Pups | 15 | 7.467 | 3.003 | 1.000 | 0.000 | 33.000 |
|  | T without Pups | 27 | 10.926 | 2.023 | 8.000 | 0.000 | 42.000 |
| **USVs and Season** | C Spring | 21 | 6.643 | 2.189 | 1.000 | 0.000 | 33.000 |
|  | T Spring | 27 | 13.296 | 2.748 | 8.000 | 0.000 | 51.000 |
|  | C Fall | 15 | 3.200 | 1.448 | 0.000 | 0.000 | 18.000 |
|  | T Fall | 15 | 6.333 | 1.866 | 5.000 | 0.000 | 24.000 |
| **Body Mass** | C | 36 | 39.210 | 0.653 | 40.000 | 30.500 | 43.000 |
|  | T | 42 | 38.500 | 0.432 | 38.000 | 34.000 | 45.000 |
| **Injection Night** | C | 36 | 3.333 | 0.080 | 3.000 | 3.000 | 4.000 |
|  | T | 42 | 3.000 | 0.000 | 3.000 | 3.000 | 3.000 |
| **USVs and Recording Night** | C - Night One | 12 | 6.750 | 2.834 | 1.500 | 0.000 | 31.000 |
|  | T - Night One | 12 | 15.071 | 3.081 | 12.000 | 0.000 | 51.000 |
|  | C - Night Two | 12 | 5.916 | 0.933 | 1.000 | 0.000 | 33.000 |
|  | T - Night Two | 14 | 9.857 | 3.959 | 9.500 | 0.000 | 42.000 |
|  | C - Night Three | 14 | 2.583 | 2.840 | 1.500 | 0.000 | 10.000 |
|  | T - Night Three | 14 | 7.500 | 3.108 | 5.000 | 0.000 | 45.000 |

**Supplemental Figure 1.**
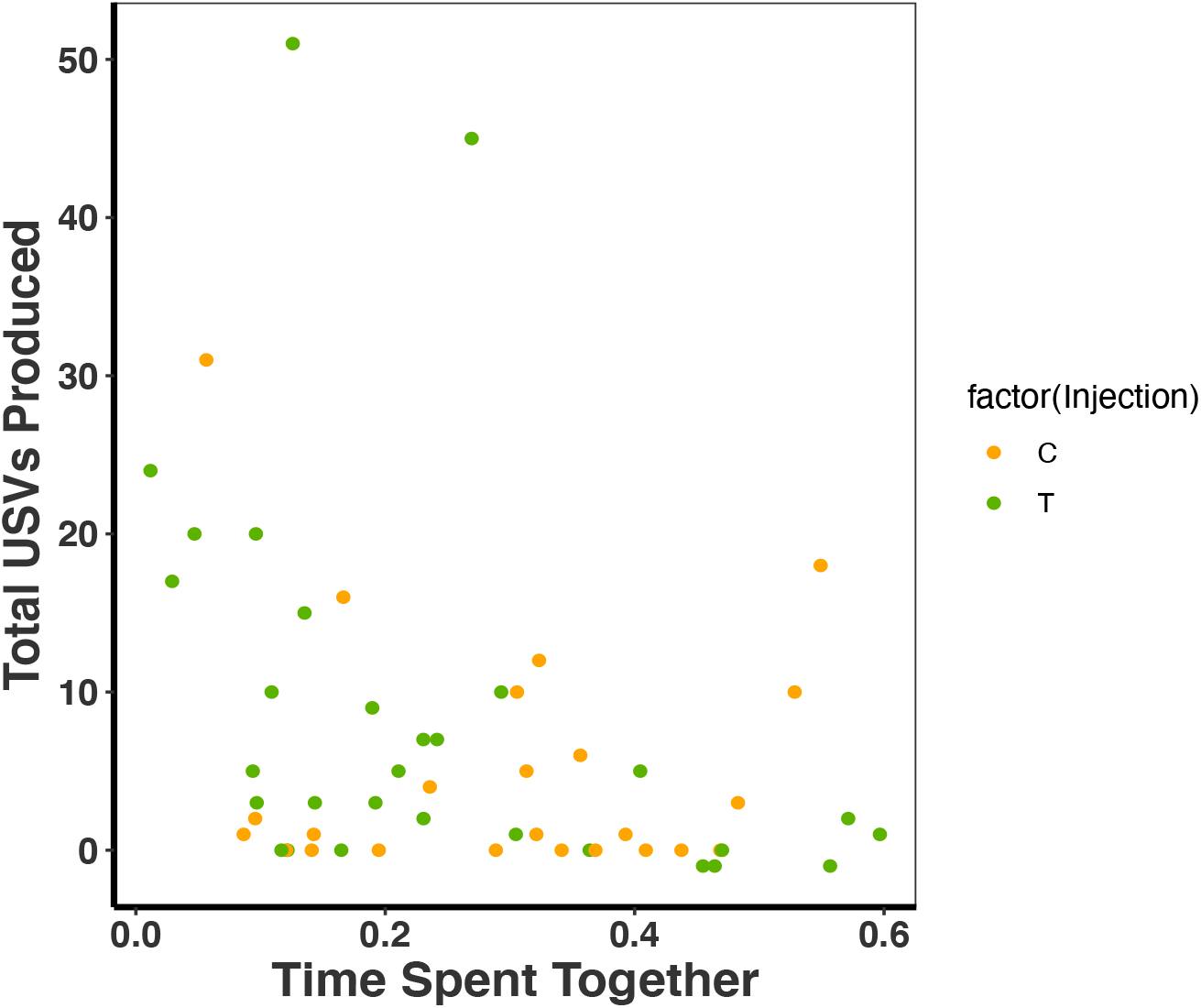
Scatterplot of the Total USVs Produced by California Mouse by Time Spent Together by the Pair at the Nest. Each female was paired with a male who received three testosterone (T= 10) or saline (C=8) injection at the nest, after the final inject, we recorded the number of USV produced and time spent at the nest for three consecutive nights. There was an indirect treatment effect on USV productions, the number of USVs produced varied by the time the pair spent together, with pairs that spent less time together produced more USVs (time spent together F_2,51_ = 20.68, R^2^= 0.12, p = 0.03; treatment = 0.37).

Both C- and T-pairs produced twice as many USVs on recording night one than on recording night three. Total USVs recorded was not influenced by pups (GLMM Estimate −0.48±0.40, p=0.25), season (GLMM Estimate −0.68±0.40, p = 0.10), body mass (GLMM Estimate 0.01±0.06, p=0.92) or total nights needed to administer all three injections (GLMM Estimate −0.85±0.64, p=0.20; Supplemental Table 4).

We have evidence that T treatment influences specific USVs directly. Regardless of distance between mates, T-pairs produced proportionately more 4SVs at the nest than control pairs (W=43, p=0.03; Supplemental Table 5). All call types (1-, 2-, 3-, 4-, 5-, 6SV, and barks) were recorded for the male and the female at both C- and T-nests. There was no significant difference between treatments in any other call type produced (1-, 2-, 3-, 5-, or 6SV; p>0.137). T-mice were more likely to call when the mate (or any other individuals besides the potential presence of pups) was not at the nest (Treatment GLM Estimate 0.72±0.11 p<0.01; mouse alone GLM Estimate 0.52±0.12 p<0.01) and there was a nonsignificant trend for more USVs produced when the mouse was far (GLM Estimate 0.22±0.13, p = 0.09). When alone (regardless of pup presence), T-mice were more likely to produce 1-, 2-, and 4SVs (1SVχ2=9.95, df=2, p<0.01; 2SVχ2 = 9.59, df=2, p<0.01; 4SVχ2 = 9.48, df=2, p<0.01). In C-mice there was no significant difference in the type of call produced (1-5SVs) when alone, or when they were close or far from another mouse (χ2=14.66, df=10, p>0.15).

**Supplemental Table 5.** Descriptive Statistics and Results from the Wilcoxon Rank Sum Test for the Comparison of USV Proportion by Type and Treatment Produced at the Nest. Each *Peromyscus californicus* male received three T (n=14) or C (n=12) injections at the nest, after the final inject, we recorded USVs at the nest for three consecutive nights. Alpha values of p <0.05 are in **bold**.

|  | USV Proportions at the Nest |  |  |  |  |  |  |  |  |
| --- | --- | --- | --- | --- | --- | --- | --- | --- | --- |
| C |  |  |  |  |  |  |  |  |  |
| Call Type | n | Mean | SE | SD | Median | Min | Max |  |  |
| 1SV | 12 | 0.205 | 0.086 | 0.299 | 0.078 | 0 | 1 |  |  |
| 2SV | 12 | 0.149 | 0.064 | 0.221 | 0.019 | 0 | 0.667 |  |  |
| 3SV | 12 | 0.251 | 0.094 | 0.326 | 0.158 | 0 | 1 |  |  |
| 4SV | 12 | 0.058 | 0.033 | 0.114 | 0 | 0 | 0.385 |  |  |
| 5SV | 12 | 0.029 | 0.019 | 0.065 | 0 | 0 | 0.194 |  |  |
| 6SV | 12 | 0.003 | 0.003 | 0.009 | 0 | 0 | 0.032 |  |  |
| BARK | 12 | 0.055 | 0.052 | 0.18 | 0 | 0 | 0.625 |  |  |
| T |  |  |  |  |  |  |  |  |  |
| Call Type | n | Mean | SE | SD | Median | Min | Max | W | p |
| 1SV | 14 | 0.287 | 0.078 | 0.293 | 0.202 | 0.029 | 1 | 60 | 0.215 |
| 2SV | 14 | 0.215 | 0.037 | 0.138 | 0.225 | 0 | 0.467 | 55 | 0.137 |
| 3SV | 14 | 0.214 | 0.045 | 0.168 | 0.2 | 0 | 0.458 | 74 | 0.602 |
| 4SV | 14 | 0.127 | 0.024 | 0.089 | 0.137 | 0 | 0.257 | 43 | 0.03 |
| 5SV | 14 | 0.014 | 0.005 | 0.018 | 0 | 0 | 0.047 | 73 | 0.525 |
| 6SV | 14 | 0.005 | 0.004 | 0.014 | 0 | 0 | 0.043 | 79 | 0.407 |
| BARK | 14 | 0.139 | 0.072 | 0.268 | 0 | 0 | 0.926 | 61 | 0.157 |

There was a direct treatment effect on call bandwidth, T-males produced calls with a 11.25% smaller bandwidth than C-males (GLM Estimate −580.22±182.47, p<0.01; Figure 2; Supplemental Table 6).

**Supplemental Table 6.** Descriptive statistics on spectral characteristics of male calls. The first call in the sequence for 1-, 2-, 3- and 4SVs produced by males (Testosterone = 86 and Control=3l).

| Acoustic Variable | Injection | n | Mean | SE | Median | Min | Max |
| --- | --- | --- | --- | --- | --- | --- | --- |
| Duration (s) | C | 31 | 0.22 | 0.02 | 0.19 | 0.06 | 0.4 |
|  | T | 86 | 0.21 | 0.01 | 0.19 | 0.06 | 0.58 |
| Bandwidth (Hz) | C | 31 | 3741.94 | 104.81 | 3900 | 2900 | 5800 |
|  | T | 86 | 3320.93 | 87.64 | 3400 | 900 | 4300 |
| Start Frequency (Hz) | C | 31 | 20367.74 | 862.22 | 19000 | 13100 | 32200 |
|  | T | 86 | 21831.4 | 465.27 | 20700 | 12600 | 33600 |
| End Frequency (Hz) | C | 31 | 21154.84 | 745.33 | 20500 | 14600 | 29700 |
|  | T | 86 | 23348.84 | 378.17 | 23400 | 15100 | 31200 |
| Frequency at Maximum Amplitude (Hz) | C | 31 | 19906.45 | 725.98 | 20500 | 13600 | 27300 |
|  | T | 86 | 22487.21 | 440.75 | 22150 | 14100 | 31700 |
| Minimum Frequency (Hz) | C | 31 | 18709.68 | 740.32 | 19500 | 12200 | 26800 |
|  | T | 86 | 21263.95 | 437.76 | 20900 | 13100 | 30200 |
| Maximum Frequency (Hz) | C | 31 | 22412.9 | 741.15 | 22900 | 16100 | 31700 |
|  | T | 86 | 24830.23 | 439.13 | 24400 | 16600 | 34100 |
| PC1 | C | 31 | -1.46 | 0.35 | -1.61 | -4.41 | 2.94 |
|  | T | 86 | -0.38 | 0.2 | -0.57 | -4.37 | 3.93 |

There was no significant difference between treatment types in call duration (GLM Estimate −0.09±0.12, p=0.46) or PC1 score (GLM Estimate 0.77±1.07, p=0.48; Supplemental Table 6). For females, there was no significant difference between treatment types and any call characteristics, duration (GLM Estimate −0.09±0.21, p=0.68), bandwidth (GLM Estimate −0.11±0.07, p=0.88) or PC1 score (GLM Estimate 0.51±1.02, p=0.63; Supplemental Table 6). There was a negative correlation between the number of USVs produced and female time at the nest (t=-1.96, df=64, p=0.05).

## Discussion

A long-standing question in the field of behavioral neuroendocrinology is what are the functions of short-term T-pulses that are induced by aggressive and sexual interactions (e.g., Ball and Balthazart 2020)? For the first time, we used a modified classic CPP paradigm to show that multiple T-pulses experienced in a location can increase the amount of time that males spend at that location. Importantly, this change in time allocation occurred in the naturally complex environment at the Hastings Natural History Reservation. We found evidence supporting the concept that the weak conditioning effects of T pulses via CPPs increased time allocation by a mammal to a location, the nest, within a territory in the wild. Our research significantly extends our understanding of the functions of such T-pulses with relevance to the large variety of species that express these T pulses. Research on California mice has provided evidence for a variety of functions of T-pulses in male California mice (review by Marler and Trainor 2020), but this is among the first study to extend this to a complex field environment (also see Timonin et al. 2018). Our research further extends evidence that T manipulations alter vocal communication, in a biparental and monogamous species through both direct and indirect mechanisms.

### Testosterone and Conditioned Place Preferences

We mimicked the natural T-pulses that occur after male-male and male-female interactions in male California mice (Marler et al. 2005; Oyegbile and Marler 2005; Zhao and Marler unpublished), as well as a number of other species (recent reviews by Maney et al 2020; Moore et al. 2020; Wingfield et al. 2020), by using T-injections. In the context of CPPs, we previously found that these injections in the laboratory can alter both time spent in a location (Zhao and Marler 2014; 2016) and social behaviors (Fuxjager et al. 2011; Pultorak et al. 2015; Timonin et al. 2018; Trainor et al. 2004; Zhao and Marler 2014; Zhao et al. 2019; 2020). Specifically, we showed that T-pulses induced that same effect in the wild at the nest site; paired male California mice experiencing three T-injections over three days spent more time at the nest and the conditioning was additive at the location in the presence of offspring. It is unlikely that the increased time at the nest was caused by behavioral changes other than the T-induced CPPs found in the laboratory for several reasons discussed below. This effect may be unique to T-pulses as compared to manipulations of T-implants used to induce long term behavioral changes in the field (e.g. Fusani 2008; Goymann et al. 2015; Ketterson et al. 1992; Marler and Moore 1989) and, among other possibilities, may allow males to avoid the high costs of sustained T levels. T-pulse release in response to social interactions can occur in a variety of species, including humans (Gleason et al. 2009; Fuxjager et al. 2017; review by Marler and Trainor 2020) and our results are consistent with laboratory observations in mice, rats, and hamsters showing that T-pulses have reinforcing/rewarding effects as described in the introduction (Alexander et al. 1994; Arnedo et al. 2000; Wood 2004; Zhao and Marler 2014; 2016). It is of interest to note that the androgen-induced CPPs can by blocked by dopamine antagonists, further reinforcing the concept of reinforcing/reward functions (Gleason et al. 2009; Marler et al. 2005; Packard et al. 1998).

T-pulses in response to male-male social challenges is a defining hallmark of Wingfield’s Challenge Hypothesis (Wingfield et al. 1990) but also occur in males after male-female sexual interaction (Gleason et al. 2009). The importance of the male-female interaction in eliciting T-pulses across species has been highlighted by Goymann (2019). Male mice and rats exposed to an estrous female or her olfactory cues show a preference for the location at which the sexual encounter occurred (Camacho et al. 2004; Frye et al. 2001; Hughes et al. 1990; Mehrara and Baum 1990). This likely serves a reproductive function as the male may use previous experiences to increase the likelihood of encounters with an estrous female and potential mating opportunity (Gleason et al. 2009). Based on the knowledge of functions of T, one might predict that increased T causes males to allocate more time toward mate guarding, courting, or aggressively pursuing other males. In the current study, the change in spatial preference was most likely not a result of behavioral changes other than the T-induced CPPs. We found no evidence for increased mate guarding behavior since females spent more time away from the nest when males spent more time at the nest. Males were not increasing their sexual behavior by pursuing their mate or other females; T-pairs spent more time apart at the nest than C-pairs. Additionally, T-males did not increase USVs associated with courtship (sweeps) that unpaired males express at high levels towards unfamiliar females (sweeps; Pultorak et al. 2015), as would be expected from courting an unfamiliar female (although these are more difficult to detect in our field set-up). This lack of increased sexual behavior to unfamiliar females is also consistent with the finding that the administration of a single T-pulse caused paired but not unpaired male California mice to decrease sweep USVs to unfamiliar females in the laboratory (Pultorak et al. 2015), suggesting an internal mate-fidelity mechanism. In the context of the nest site, there was no evidence that T-pulses increased aggression (see laboratory studies focused on male-male interactions; Trainor et al. 2004), as evidenced by a lack of injuries (all animals tested were trapped post experiment with no visible injuries) or increase in aggressive barks (Pultorak et al. 2018). We cannot, however, rule out that males may have been actively pushing females out of the nest as has been anecdotally observed in laboratory situations by either sex (Rieger and Marler, unpublished data). What then were males doing at the nest? In this case, the most likely explanation is increased paternal behavior (when pups are at the nest) in the form of increased nest defense or paternal care of pups based on evidence, described below, that T can increase paternal care in California mice in the laboratory as a by-product of spending more time at the nest. We suggest that T increases the focus on the reproductive or aggressive behaviors most relevant at that time depending on the social and physical contexts for that specific species (Hurley and Kalcounis-Rueppell 2018). This is consistent with previous findings that the ability to create T-induced conditioned location preferences is plastic and varies with social experience and current social and physical (e.g. familiar versus unfamiliar locations) contexts (Zhao and Marler 2016). It would be valuable in the future to examine the natural expression of T-pulses in males in response to social stimuli in the field. We cannot rule out the alternative that males simply spend more time at the nest without altering paternal or direct pup defense behaviors.

In nature, T-pulse release following a sexual encounter most likely occurs at the nest site when females first approach a male that has established a territory, further T-pulses likely occur when the female is in postpartum estrous (Gubernick and Nelson 1989) and T-induced CPPs could be the potential mechanism for increasing paternal care through increased preference for spending time at the nest. Therefore, T-induced CPPs at the nest could drive the male to spend more time at the nest where future sexual encounters are more likely to take place, which could also be a mechanism for maintaining paternal care. In addition, in California mice and other species, T promotes paternal care in males (for example, Juana et al. 2009; Trainor and Marler 2002; Ziegler et al. 2004). California mouse pups demand extensive paternal investment because they are altricial and exothermic and depend on adult presence to maintain their body temperature (Gubernick and Alberts 1987). In the California mouse, the presence of the father has a significant positive effect on offspring survival when temperatures are low and the parents have to forage, but there is no effect of father’s presence on pup survival when exposed to warm temperatures in the laboratory (Gubernick et al. 1993). The importance of the father, however, is highlighted by findings in the wild that paternal presence has a significant positive effect on offspring survival in the field (Gubernick and Teferi 2000), as well as in laboratory studies (Bambico et al. 2013; Cantoni and Brown 1997; Rosenfeld et al. 2013). The main limiting factor in California mouse reproduction is water availability (Nelson et al. 1995) and therefore, in nature, California mice breed during the cold rainy season and cease breeding during the dry summer months (Nelson et al. 1995). When reproduction occurs during harsh environmental conditions and offspring require constant care, there must be a balance in the time invested towards offspring maintenance and time spent towards foraging and resource defense. To achieve balance, biparental care is essential for facilitating offspring survival and maximizing reproductive success. We, therefore, propose that in some biparental species, T-induced CPPs could be a mechanism for keeping the male at the nest to care for the young while the female forages or conducts other behaviors related to territory maintenance. Another selection pressure for T-induced paternal behavior may be increased protectiveness of pups to prevent the high levels of conspecific infanticide found in rodents (Agrell et al. 1998). Van Anders et al. (2012) speculate that infant protection may be positively associated with T and more nurturing behaviors negatively associated with T. The reinforcing effects of T-pulses may function to allocate more time in the familiar environment and display behaviors that have direct fitness benefits.

Interestingly, changes in male allocation to tasks closer to the nest resulted in females spending more time away from the nest. This could occur through female preference/choice or because males aggressively pushed females out of the nest. We again do not think that there was an increase in aggression between the mated pair because of the absence of increased barks at the nest as has been shown in mated pairs that are stressed (such as through a fidelity challenge; Pultorak et al. 2018). An alternative possibility is that females change their spatial preference to be away from the nest to compensate for the T-induced changes in male spatial preferences. We also observed plasticity in female but not male time at the nest in different seasons, suggesting plasticity in maternal behavior in response to environmental factors. In this study we speculate that plasticity in the males is influenced by T from social stimuli, whereas the plasticity we see in the females may be influenced more directly by the physical environment. In species that form pair-bonds where both members of a pair are engaged in offspring care and territory defense, the delegation of tasks is beneficial. In a wider variety of taxonomic groups, including insects, birds, fish, and mammals that engage in cooperative breeding, members of a pair or group often distribute tasks (Arnold et al. 2005; Ahern, et al. 2011; Mathews 2002; Page et al. 2006; Quinard and Cézilly 2012; Rieger et al. 2019; Rogers 1988). In the laboratory, when challenged with a potential intruder, California mouse pairs either coordinate their behavior in joint defense or employ labor division strategies, with the latter strategy potentially more likely to occur after pups are born (Rieger et al. 2019). In the California mouse, when the male is present but decreases paternal care due to castration, the female compensates for the mate’s behavior by increasing the huddling of her pups (Trainor and Marler 2001). Species in which both members provide offspring care, such as in the Midas cichlid, great tit, and prairie vole, the presence of offspring increases the pairs’ use of division of labor (Ahern et al. 2011; Boucaud et al. 2016; Rogers et al. 2018; Rogers 1988). This division of labor can have important long-term benefits for the persistence and survival of a social group (Arnold et al. 2005). In the case of California mice, if the male is spending more time in one location, such as the nest to care for offspring, then the female is adjusting her space use by allocating more time to other parts of the territory, such as foraging and/or defending the territory against potential intruders. Interestingly, T-induced CPPs regulate behaviors of other animals by influencing space use which in turn alters social interactions.

### Testosterone and Vocal Communication

We also found that the same transient increases in T which induced CPPs also had long-term effects (>24 hours) on vocal communication by increasing the number of USVs produced and altering both the type of calls produced and the call bandwidth. Our results are consistent with Timonin et al (2018) who examined the effect of T pulses on USVs in this species in the wild; T-pairs from both studies produced more total calls across the three nights with fewer USVs produced on night three. One difference between the studies is that Timonin et al (2018) found that T-pairs produced proportionately more 1-, 4- and 5SVs, whereas we only found a difference in 4SVs; we also found, however, that T-pairs were more likely to produce 1-, 2-, and 4SVs when alone. The difference between the studies could be attributed to year, population densities, or a higher sample size in the current study. Anecdotally, densities were lower in the current study which could alter social interactions. The current study also reveals that the increased time apart in T-pairs may indirectly drive the greater number of USVs produced by the T-pairs. However, while pairs call more when separated, T significantly increased calling rate when a member of a pair was alone, with a nonsignificant trend when pairs were far apart (p = 0.09).

We further discovered that the increase in SV production was associated with a decrease in bandwidth. An intriguing speculation is that the narrower bandwidth USVs produced are less susceptible to environmental degradation and may travel further in the environment (Barber et al. 2010; Slabbekoorn 2013; Zhang et al. 2015). The narrow bandwidth SVs may therefore be more efficient at reaching the females when males and females are spending more time apart in response to the T-manipulation. Contrary to our findings that T-pulses decreased bandwidth in SVs, in the golden hamsters (*Mesocricetus auratus*) T-pulses increased bandwidth of the one and multi-note sudden frequency change calls, however, these were produced in a sexual context in close proximity (Fernández-Vargas 2017). Furthermore, males in singing mice (*Scotinomys teguina*) that are administered T-implants produce frequency modulated trills (mating calls) also with increased bandwidth, which females tended to prefer (Pasch et al. 2011a; Pasch et al. 2011b). We speculate that under the conditions of male-female interactions in a mate-choice context, the function of the bandwidth change may be related to the increased call complexity and greater information transfer characteristic of wider bandwidths. In birds and some frogs, however, while narrow bandwidth calls are less attractive to novel females, these calls can also transmit more effectively in the soundscape (Francis et al. 2011; Montague et al. 2013; Slabbekoorn 2013). We suspect that California mice do not follow the same pattern of call production as golden hamsters and singing mice because monogamous California mouse calls in the current study are likely directed toward the other member of the already established pair located elsewhere on the territory (Briggs and Kalcounis-Rueppell 2011). Moreover, it is unlikely to be directed towards pups because in the current study offspring presence did not influence call production. In the wild, SVs recorded from unmanipulated California mice traveled an average of 3.12 m and a maximum of 7 m (Timonin et al. 2018). Narrow bandwidth calls provide less information but increases the likelihood of signal transmission of the vocal signal across a large area (Barber et al. 2010). When males alter bandwidth of SV calls, it is unlikely that the call functions to attract a new mate, but instead likely elicits attention of his own mate and maintains awareness of the other individual’s location as suggested by Briggs and Kalcounis-Rueppell (2011). The increased calling behavior when the pair is apart but decreased calling behavior when the pair is together could be a contact call or function to reduce extra-pair copulation and maintain sexual fidelity. Again, a single T injection also decreased a paired male’s USV response to an unfamiliar female, suggesting that T pulses in pairs of this monogamous species do not function to attract a male to a new potential mate through USVs, in fact, simple sweeps are likely more important in courtship that SVs (Pultorak et al. 2015; Pultorak et al. 2018). It is also possible, however, and remains untested, that the calls serve a dual function, as mate contact calls and/or as territorial advertisement. Our data suggest that call production most likely serves for maintaining awareness of the other individuals in a complex environment (Hurley and Kalcounis-Rueppell 2018).

In summary, this is the first field study that demonstrates a potentially natural function of transient T-pulses, that of inducing CPPs. T-pulses naturally occur in a variety of different species, including humans (Fuxjager et al. 2017), and our results are consistent with other research in which T-pulses have rewarding properties and can condition animals to the physical location in which the hormone release occurred (e.g. Arnedo et al. 2000; Frye et al. 2001). We now know that despite T being weakly reinforcing compared to many drugs, it can alter behavior and do so in a complex natural environment. This change in the allocation of time spent in specific physical environments also leads to changes in call production during those interactions, likely resulting, in part, from T-induced changes in social interactions. In this case, specifically altering male time spent at the nest, with the possibility of an increase in paternal behavior, and with a complimentary change in the unmanipulated mate. We speculate that there could be an adaptive significance for a co-option mechanism that allows a close association between mating release of T and paternal behavior. While we have effectively demonstrated potential functions of T-pulses in the laboratory and field through the current and previous studies, we do not yet know if these functions differ from those of T-implants that mimic the longer lasting seasonal changes such as breeding versus nonbreeding season (Wingfield et al. 2000). We speculate, however, that the T-pulses are tied in with active learning from a changing social environment during the breeding season in relation to functions related to reproduction. Once thought to be of little importance, especially in humans (Geniole et al. 2020), we are discovering that T-pulses have the potential to allow males to adjust to changing social conditions in the wild through both spatial preference and vocal plasticity of a male and his mate.

## Methods

Field work was conducted at the Hastings Natural History Reservation (HNHR), Carmel Valley, California, USA, from January to June 2015 (season one) and from September to December 2015 (season two) on established trapping grids (see Briggs and Kalcounis-Rueppell 2011; Kalcounis-Rüppell and Millar 2002; Kalcounis-Rueppell et al. 2006, 2010; Timonin et al. 2018;). We tagged 323 mice, of those we identified 33 reproductively active mated pairs (males with enlarged testis and females were pregnant and/or lactating). Once putative pairs were identified, we trapped the target and both the male and the female in the pair were outfitted with a 0.55g M1450 mouse style transmitter (Advanced Telemetry System [ATS], Isanti, MN, USA), adjusted for California mice (Briggs and Kalcounis-Rueppell 2011). We attached the transmitters (Briggs and Kalcounis-Rueppell 2011) and released all mice at the site of capture. Using an R4500S DCC receiver/datalogger and a Yagi antenna (ATS), we located the pair the following day at the nest (described below). All 33 putative pairs were confirmed as pairs when the signals from both the male and female transmitters were emitted from the same nest. We ensured that the tracked nest location was the primary nest and not one of the satellite locations by monitoring nest occupancy for up to three days. A total of 28 pairs were in the nest for up to three days post-tracking, and we ensured that the nest was in a suitable location for setting-up our remote sensing equipment (described below). We placed 15-20 Longworth traps (14 x 6.5 x 8.5cm, NHBS, Totnes, Devon, UK) within a 2m radius surrounding the nest to trap the male and administer injections (described below).

### Treatment

We randomly assigned 28 males to receive either T (n=15) or saline (control, C, n=13) injections. The dose of T injection was approximately 36ug/kg (T-cyclodextrin dissolved in saline) which mimics natural T-pulses (Oyegbile and Marler 2005; Trainor et al. 2004) and has been used successfully in multiple studies with male California mice primarily focused on aggression but including courtship (Fuxjager et al. 2011; Pultorak et al. 2015; Timonin et al. 2018; Trainor et al. 2004; Zhao and Marler 2014; Zhao et al. 2020; 2019;). All animals were injected subcutaneously, and the researcher was blind to the treatment type. Each focal male received three injections of 0.1 ml of the injectate regardless of body mass. We, therefore, included body mass as an independent variable in our statistical analysis. All three injections were administered within five days, with only one injection per day. One male was excluded because he did not receive all three injections within five days. We refer to females whose mate received T as “T-females” and the nests as “T-nests”. Females whose mate received saline are referred to as “C-females” and the nests as “C-nests”. We also recorded total number of nights needed to administer all three injections (three or four nights), therefore, we included total nights as an independent variable in our statistical analysis. After the third and last injection, we deployed the remote sensing equipment (automated radio telemetry, audio recording, and thermal imaging; described below) to record for three consecutive nights (“recordings nights” 1-3). We treated data collected by the remote sensing equipment over one night as a sample unit and included recording night in our analyses. For each recording session, all equipment was set-up to record from sunset to sunrise. T and C solutions were provided by Dr. Brian Trainor from the Department of Psychology at the University of California Davis (IACUC Protocol number 19849).

### Automated Radio Telemetry

We used two R4500S DCC receiver/dataloggers (Advanced Telemetry System [ATS], Isanti, MN, USA) to monitor the number of minutes radio-collared mice spent at the nest each night and the amount of time the male and female were together and apart. Each datalogger was connected to an antenna and programmed to detect one unique transmitter frequency (one for each pair member). Antennas were placed either on top of or next to the nest. When the collared mouse was detected by the receiver, signal strength was stored in the datalogger. We, therefore, monitored both male behavioral changes in response to treatment type and the female response to male behavioral changes. The time at the nest was standardized, because of differences in length of recordings, by counting and totaling the number of minutes the mouse spent in the nest area from sunset to sunrise divided by the total number of minutes in the night to obtain a proportion of time spent in the nest area (“time at the nest”). Each day we also conducted manual telemetry on the collared pair and found the nest location with the strongest signal strength. For each individual, we assessed a reference signal (range 130 – 155dB signal strength) during the day when we knew the mouse was in the nest. To assess how long a mouse spent in the nest per night, we only counted the number of minutes during which the signal fell within the reference range. Each morning, the dataloggers were removed from the field and data were downloaded. The telemetry equipment was set-up at 27 nest sites. Due to equipment failure, we did not record male time at the nest for five T-nests and one C-nest and we did not record female time at the nest for one T-nest and three C-nests. Our final dataset consisted of 63 recording nights from 21 nest sites (T=10, C=11) for males and 69 recording nights from 23 nest sites (T=14, C=9) for females. We did not have matching pair time at the nest for five T-nests and four C-nests. Our final matching pair dataset consisted of 54 recording nights from 18 nest sites (T=10, C=8) and we used night as a sample unit in our analysis.

### Audio Recording

Our goal was to record a variety of USVs as described here. The SVs have a peak frequency around 20kHz, and are approximately 50 – 1000ms in length, low modulation calls that can be emitted as a single or bout of multiple calls that can be categorized based on the number of calls in a bout (1SV, 2SV, 3SV, 4SV, etc.; Kalcounis-Rueppell et al. 2018). Bark calls are shorter in duration (50ms or less), resemble an upside-down U with the beginning and the end of the call dips into audible range at approximately 12kHz with a peak frequency around 20kHz and tend to be “noisy” vocalizations (Pultorak et al. 2018). Complex sweeps are short duration (100ms or less), that pass through multiple high to low and low to high frequencies with a peak frequency of around 100kHz (Pultorak et al. 2015). Similar to the SVs, the barks, simple sweeps, and complex sweeps occur as a single call or bout of calls.

We used ultrasonic microphones (Emkay FG Series from Avisoft Bioacoustics, Berlin, Germany) to assess the number and type of USVs produced at the nest. We set up two microphones; one next to the nest entrance and a second 2m away directly from the nest entrance. Microphones recorded as described in Timonin et al. 2018. When possible, we assigned USVs to individuals by matching the radio telemetry data with the time of the mouse USV. By examining telemetry data within one minute of USV production and based on the transmitter signal strength (Briggs and Kalcounis-Rueppell 2011), we determined if the male or the female produced the USV. We were not able to assign 51% of the USVs to one individual because both the male and the female were at the nest with strong transmitter signal strengths and therefore, we only used the assigned data to test the treatment effect on the spectral and temporal characteristics of USVs. The acoustic recording system was set-up at 27 nest sites (T=15, C=12). Due to equipment failure, we did not record data at one T-nest. Our final dataset consisted of 78 recording nights from 26 nest sites (T=14, C=12). Mouse USVs were counted and classified into one of the following types: 1SV, 2SV, 3SV, 4SV, 5SV, 6SVs or barks (Kalcounis-Rueppell et al. 2018). We counted USV numbers recorded from sunset to sunrise and refer to the value as “total USVs”. Lastly, we determined if the proportion of a specific type of USV (1-, 2-, 3-, 4-, 5-, 6SVs and barks) differed between treatments by totaling each USV type per nest site and dividing by the total number of USVs produced at that nest.

Using SAS Lab Pro, we extracted spectral and temporal characteristics (duration, bandwidth [number of frequencies a call passes through], and five frequency variables [peak, minimum, maximum, start, and end]) from USVs recorded at the nest. Each spectrogram was generated with a 512 FFT (Fast Fourier Transform), and a 100-frame size with Hamming window. For each call, we measured duration, bandwidth, and five frequency parameters (start, end, minimum, maximum, and frequency at maximum amplitude).

### Thermal Imaging

We used a thermal imaging lens (Photon 320 14.25 mm; Flir/Core By Indigo) to assign social context to USVs. The thermal imaging lens was suspended to capture the full view of the nest and a circular area with a 2m radius surrounding the nest. The lens was connected to a JVC Everio HDD camcorder which recorded continuously throughout the night. We watched the video footage in three-minute increments, (1-minute before, 1-minute during and 1-minute after call production) to determine behavior and number of mice on the screen. If there was only one member of a pair present at a time, the behavior was assigned as “alone”. If both mates were present, we determined the proximity of mice to each other by using a 1m scale that was overlaid in the video for each site. If mice were less than 1m apart, we assigned them as “<1m”, and if the mice were more than 1m apart, we marked them as “>1m”. We assessed the types of USVs (1-, 2-, 3-, 4-, 5-, 6SVs and barks) produced by context (alone, <1m or >1m) and treatment type.

### Statistical Analyses

Time at the nest for both the male and the female was normally distributed and therefore we fitted a Gaussian distribution. Pair time at the nest and total USVs were in violation of normality and variances and could not be normalized and therefore we used either a Quasibinomial or Poisson distribution. We used General Linear Models (GLM) with time at the nest, pair time at the nest and total USVs as the dependent variables and we included individual identification code independent of treatment type to account for individual differences. Using the package lme4 (Bates et al. 2015), we fitted Generalized Linear Mixed Models (GLMM) with the individual identification code as a random term and treatment as the fixed term.

In addition to treatment type, we also considered the following covariates: presence of pups at the nest, season, male and female body mass, total nights needed to administer all three injections, and recording night. Due to our small sample size, when modeling covariates we included a maximum of two fixed terms in one GLMM model (treatment type and one covariate). We first modeled the interaction term between treatment type and the one covariate. If the interaction term was not significant, the term was dropped. We also used the non-parametric Wilcoxon Rank Sum test for our comparison of USV types. We compared the median of the proportion of each USV type by treatment. We used a GLM to examine the relationship between USV types by context and treatment.

For the analysis of the spectral and temporal characteristics, we used factor analysis to extract principal component (PC) scores for the frequency parameters (as in Kalcounis-Rueppell et al. 2010). We only analyzed calls assigned to an individual male or female and the calls were analyzed separately. We generated a single PC score that represented the frequency variables using the first call in the 1-, 2-, 3- and 4SVs sequence. We did not include 5SVs, 6SVs, and barks due to a small sample size (<4). PC1 accounted for 67% of the variation in acoustic variables for male calls and 71% variation for female calls. We fitted GLMM with ID as a random term and USV type and treatment as the fixed terms. For both male and female calls, duration and bandwidth variables were in violation of normality and variances. We, therefore, fitted our models using a Poisson family distribution. PC scores were normally distributed, and we used a Gaussian distribution in our models. All data are represented using box plots. We used an alpha level of p<0.05 for the rejection criterion. All data were analyzed using R software (Version 3.2.2.)

## Acknowledgments

We thank A. Campos, J. Caprio, C. Falvo, M. Grupper, C. Kovarik, A. Larsen, J. Neill, and E. Sakonjic for their assistance in data collection and Brian Trainor for providing our C and T injectate. We also thank J. del Valle, V. Voegeli and Hastings Natural History Reserve for their support and providing the use of their facilities during our field season. Lastly, we appreciate the input from H. Li on all aspects of the analysis and B. Trainor, R. Bhandari, G. Wasserberg and C. Snowdon for comments on the manuscript. This work was supported by the National Science Foundation (NSF; IOS-1355163) and UNC-Greensboro.

## References

Agrell J, Wolff JO, and Ylönen H. 1998. Counter-strategies to infanticide in mammals: costs and consequences. Oikos 83 (3): 507–17. https://doi.org/10.2307/3546678.

Ahern TH, Hammock EAD, and Young LJ. 2011. Parental division of labor, coordination, and the effects of family structure on parenting in monogamous prairie voles (*Microtus ochrogaster*). Dev Psychobiol 53 (2): 118–31. https://doi-org.libproxy.uncg.edu/10.1002/dev.20498.

Alexander GM, Packard MG, and Hines M. 1994. Testosterone has rewarding affective properties in male rats: implications for the biological basis of sexual motivation. Behavioral Neuroscience 108 (2): 424–28.

Anders SM, Tolman RM, and Volling BL. 2012. Baby cries and nurturance affect testosterone in men. Hormones and Behavior 61 (1): 31–36. https://doi.org/10.1016/j.yhbeh.2011.09.012.

Arnedo MT, Salvador A, Martinez-Sanchis S, and Gonzalez-Bono E. 2000. Rewarding properties of testosterone in intact male mice: a pilot study. Pharmacology Biochemistry and Behavior 65 (2): 327–32. https://doi.org/10.1016/S0091-3057(99)00189-6.

Arnold KE, Owens IF, and Goldizen AW. 2005. Division of labour within cooperatively breeding groups. Behaviour 142 (11–12): 1577–90. https://doi.org/10.1163/156853905774831927.

Bambico FR, Lacoste B, Hattan PR, and Gobbi G. 2013. Father absence in the monogamous california mouse impairs social behavior and modifies dopamine and glutamate synapses in the medial prefrontal cortex. Cerebral Cortex 25 (5): 1163–75. https://doi.org/10.1093/cercor/bht310.

Ball GF and Balthazart J. 2020. The neuroendocrine integration of environmental information, the regulation and action of testosterone and the challenge hypothesis. Hormones and Behavior Jul;123:104574. doi: 10.1016/j.yhbeh.2019.104574.

Barber JR, Crooks KR, and Fristrup KM. 2010. The costs of chronic noise exposure for terrestrial organisms. Trends in Ecology and Evolution 25 (3): 180–89. https://doi.org/10.1016/j.tree.2009.08.002.

Bates D, Mächler M, Bolker B, and Walker S. 2015. Fitting linear mixed-effects models using Lme4. Journal of Statistical Software 67 (1): 1–48. https://doi.org/10.18637/jss.v067.i01.

Bell MR, and Sisk CL. 2013. Dopamine mediates testosterone-induced social reward in male Syrian hamsters. Endocrinology 154 (3): 1225–34. https://doi.org/10.1210/en.2012-2042.

Boucaud ICA, Aguirre Smith MLN, Valère PA, and Vignal C. 2016. Incubating females signal their needs during intrapair vocal communication at the nest: a feeding experiment in great tits. Animal Behaviour 122: 77–86. https://doi.org/10.1016/j.anbehav.2016.09.021.

Briggs JR, and Kalcounis-Rueppell MC. 2011. Similar acoustic structure and behavioural context of vocalizations produced by male and female California mice in the wild. Animal Behaviour 82 (6): 1263–73. https://doi.org/10.1016/j.anbehav.2011.09.003.

Camacho F, Sandoval C, and Paredes R. 2004. Sexual experience and conditioned place preference in male rats. Pharmacology Biochemistry and Behavior, Sex and Drugs, 78 (3): 419–25. https://doi.org/10.1016/j.pbb.2004.04.015.

Cantoni D, and Brown RE. 1997. Paternal investment and reproductive success in the California mouse, *Peromyscus californicus*. Animal Behaviour 54 (2): 377–86. https://doi.org/10.1006/anbe.1996.0583.

Fernández-Vargas M. 2017. Rapid effects of estrogens and androgens on temporal and spectral features in ultrasonic vocalizations. Hormones and Behavior 94: 69–83. https://doi.org/10.1016/j.yhbeh.2017.06.010.

Francis CD, Ortega CP, and Cruz A 2011. Vocal frequency change reflects different responses to anthropogenic noise in two suboscine tyrant flycatchers. Proceedings of the Royal Society B: Biological Sciences, 278, 2025–2031

Frye CA, Park D, Tanaka M, Rosellini R, and Svare B. 2001. The testosterone metabolite and neurosteroid 3α-androstanediol may mediate the effects of testosterone on conditioned place preference. Psychoneuroendocrinology 26 (7): 731–50. https://doi.org/10.1016/S0306-4530(01)00027-0.

Fusani L. 2008. Endocrinology in field studies: problems and solutions for the experimental design. General and Comparative Endocrinology 157 (3): 249–53. https://doi.org/10.1016/j.ygcen.2008.04.016.

Fuxjager MJ, Montgomery JL, and Marler CA. 2011. Species differences in the winner effect disappear in response to post-victory testosterone manipulations. Proceedings of the Royal Society B: Biological Sciences 278 (1724): 3497–3503. https://doi.org/10.1098/rspb.2011.0301.

Fuxjager MJ, Trainor BC, and Marler CA. 2017. What can animal research tell us about the link between androgens and social competition in humans? Hormones and Behavior, Hormones and Human Competition, 92: 182–89. https://doi.org/10.1016/j.yhbeh.2016.11.014.

Geniole SN, Bird BM, McVittie JS, Purcell RB, Archer J, Carré JM. 2020. Is testosterone linked to human aggression? A meta-analytic examination of the relationship between baseline, dynamic, and manipulated testosterone on human aggression. Hormones and Behavior Jul;123:104644. doi: 10.1016/j.yhbeh.2019.104644.

Gleason ED, Fuxjager MJ, Oyegbile TO, and Marler CA. 2009. Testosterone release and social context: when it occurs and why. Frontiers in Neuroendocrinology 30 (4): 460–69. https://doi.org/10.1016/j.yfrne.2009.04.009.

Glickman SE, and Schiff BB. 1967. A biological theory of reinforcement. Psychological Review 74 (2): 81–109. https://doi.org/10.1037/h0024290.

Goymann W, Moore IT, and Oliveira RF. 2019. Challenge hypothesis 2.0: a fresh look at an established idea. BioScience 69 (6): 432–42. https://doi.org/10.1093/biosci/biz041.

Goymann W, Villavicencio CP, and Apfelbeck B. 2015. Does a short-term increase in testosterone affect the intensity or persistence of territorial aggression? An approach using an individual’s hormonal reactive scope to study hormonal effects on behavior. Physiology and Behavior 149: 310–16.

Gubernick DJ, and Alberts JR. 1987. The biparental care system of the California mouse, *Peromyscus californicus*. Journal of Comparative Psychology 101 (2): 169–77. https://doi.org/10.1037/0735-7036.101.2.169.

Gubernick DJ, and Nelson RJ. 1989. Prolactin and paternal behavior in the biparental California mouse, *Peromyscus californicus*. Hormones and Behavior 23 (2): 203–10. https://doi.org/10.1016/0018-506X(89)90061-5.

Gubernick DJ, Wright SL, and Brown RE. 1993. The significance of father’s presence for offspring survival in the monogamous California mouse, *Peromyscus californicus*. Animal Behaviour 46 (3): 539–46. https://doi.org/10.1006/anbe.1993.1221.

Gubernick DJ, and Teferi T. 2000. Adaptive Significance of male parental care in a monogamous mammal. Proceedings of the Royal Society of London. Series B: Biological Sciences 267 (1439): 147–50. https://doi.org/10.1098/rspb.2000.0979.

Hughes AM, Everitt BJ, and Herbert J. 1990. Comparative effects of preoptic area infusions of opioid peptides, lesions and castration on sexual behaviour in male rats: studies of instrumental behaviour, conditioned place preference and partner preference. Psychopharmacology 102 (2): 243–56. https://doi.org/10.1007/BF02245929.

Hurley L, and Kalcounis-Rueppell M. 2018. State and context in vocal communication of rodents. In: Dent ML, Fay RR, and Popper AN, editors, Rodent Bioacoustics, Springer Handbook of Auditory Research Cham, Switzerland, Springer 67: 202–221.

Juana L, Ramírez L, Carmona A, Ortiz V, Delgado J, and Cardenas R. 2009. Paternal behavior and testosterone plasma levels in the volcano mouse *Neotomodon Alstoni* (Rodentia: muridae). Revista de Biología Tropical 57: 433–39. https://doi.org/10.15517/rbt.v57i1-2.11360.

Kalcounis-Rüppell MC, and Millar JS. 2002. Partitioning of space, food, and time by syntopic *Peromyscus Boylii* and *P. Californicus*. Journal of Mammalogy 83 (2): 614–25. https://doi.org/10.1644/1545-1542(2002)083<0614:POSFAT>2.0.CO;2.

Kalcounis-Rueppell MC, Metheny JD, and Vonhof MJ. 2006. Production of ultrasonic vocalizations by *Peromyscus* mice in the wild. Frontiers in Zoology 3: 3. https://doi.org/10.1186/1742-9994-3-3.

Kalcounis-Rueppell MC, Petric R, Briggs JR, Carney C, Marshall MM, Willse JT, Rueppell O, Ribble DO, and Crossland JP. 2010. Differences in ultrasonic vocalizations between wild and laboratory California mice (*Peromyscus californicus*). PLOS ONE 5 (4): e9705. https://doi.org/10.1371/journal.pone.0009705.

Kalcounis-Rueppell MC, Pultorak JD, and Marler CA. 2018. Chapter 22 - Ultrasonic vocalizations of mice in the genus *Peromyscus*. In Handbook of Ultrasonic Vocalization, edited by Stefan M. Brudzynski, Handbook of Behavioral Neuroscience. Elsevier 25:227–35. https://doi.org/10.1016/B978-0-12-809600-0.00022-6.

Ketterson ED, Nolan V, Wolf L, and Ziegenfus C. 1992. Testosterone and avian life histories: effects of experimentally elevated testosterone on behavior and correlates of fitness in the dark-eyed junco (*Junco hyemalis*). The American Naturalist 140 (6): 980–99.

Maney, DL, Aldredge RA, Edwards SA, James NP, and Sockman KW. 2020. “Time Course of Photo-Induced Egr-1 Expression in the Hypothalamus of a Seasonally Breeding Songbird.” Molecular and Cellular Endocrinology 512: 110854. https://doi.org/10.1016/j.mce.2020.110854.

Marler CA, and Moore MC. 1989. Time and energy costs of aggression in testosterone-implanted free-living male mountain spiny lizards (*Sceloporus jarrovi*). Physiological Zoology 62 (6): 1334–50. https://doi.org/10.1086/physzool.62.6.30156216.

Marler CA, Oyegbile TO, Plavicki J, and Trainor BC. 2005. Response to Wingfield’s commentary on ‘A continuing saga: the role of testosterone in aggression.’ Hormones and Behavior 48 (3): 256–58. https://doi.org/10.1016/j.yhbeh.2005.05.010.

Marler CA, and Trainor BC. 2020. The challenge hypothesis revisited: focus on reproductive experience and neural mechanisms. Hormones and Behavior 123: 104645. https://doi.org/10.1016/j.yhbeh.2019.104645.

Mathews LM. 2002. Territorial cooperation and social monogamy: factors affecting intersexual behaviours in pair-living snapping shrimp. Animal Behaviour 63 (4): 767–77. https://doi.org/10.1006/anbe.2001.1976.

Mehrara BJ, and Baum MJ. 1990. Naloxone disrupts the expression but not the acquisition by male rats of a conditioned place preference response for an oestrous female. Psychopharmacology 101 (1): 118–25. https://doi.org/10.1007/BF02253728.

Montague MJ, Danek-Gontard M, and Kunc HP. 2013. Phenotypic plasticity affects the response of a sexually selected trait to anthropogenic noise. Behavioral Ecology, 24, 342–348. https://doi.org/10.1093/beheco/ars169.

Moore IT, Hernandez J, and Goymann W. 2020. “Who Rises to the Challenge? Testing the Challenge Hypothesis in Fish, Amphibians, Reptiles, and Mammals.” Hormones and Behavior 123: 104537. https://doi.org/10.1016/j.yhbeh.2019.06.001.

Nelson RJ, Gubernick DJ, and Blom JMC. 1995. Influence of photoperiod, green food, and water availability on reproduction in male California mice (*Peromyscus californicus*). Physiology and Behavior 57 (6): 1175–80. https://doi.org/10.1016/0031-9384(94)00380-N.

Oyegbile TO, and CA Marler. 2005. Winning Fights Elevates Testosterone Levels in California Mice and Enhances Future Ability to Win Fights. Hormones and Behavior 48 (3): 259–67. https://doi.org/10.1016/j.yhbeh.2005.04.007.

Packard MG, Schroeder JP, Alexander GM. Expression of testosterone conditioned place preference is blocked by peripheral or intra-accumbens injection of alpha-flupenthixol. Hormones and Behavior 1998 Aug;34(1):39–47. doi: 10.1006/hbeh.1998.1461. PMID: 9735227.

Page RE, Scheiner R, Erber J, and Amdam GV. 2006. The development and evolution of division of labor and foraging specialization in a social insect (Apis mellifera l.). 7 Current Topics in Developmental Biology. Academic Press 4:253–86.https://doi.org/10.1016/S0070-2153(06)74008-X.

Pasch B, George AS, Campbell P, and Phelps SM. 2011a. Androgen-dependent male vocal performance influences female preference in neotropical singing mice. Animal Behaviour 82 (2): 177–83. https://doi.org/10.1016/j.anbehav.2011.04.018.

Pasch B, George AS., Hamlin HJ, Guillette LJ, and Phelps SM. 2011b. Androgens modulate song effort and aggression in neotropical singing mice. Hormones and Behavior 59 (1): 90–97. https://doi.org/10.1016/j.yhbeh.2010.10.011.

Pultorak JD, Fuxjager MJ, Kalcounis-Rueppell MC, and Marler CA. 2015. Male fidelity expressed through rapid testosterone suppression of ultrasonic vocalizations to novel females in the monogamous California mouse. Hormones and Behavior 70: 47–56. https://doi.org/10.1016/j.yhbeh.2015.02.003.

Pultorak JD, Matusinec KR, Miller ZK, and Marler CA. 2017. Ultrasonic vocalization production and playback predicts intrapair and extrapair social behaviour in a monogamous mouse. Animal Behaviour 125: 13–23. https://doi.org/10.1016/j.anbehav.2016.12.023.

Pultorak JD, Alger SJ, Loria SO, Johnson AM, and Marler CA. 2018. Changes in behavior and ultrasonic vocalizations during pair bonding and in response to an infidelity challenge in monogamous California mice. Frontiers in Ecology and Evolution 6. https://doi.org/10.3389/fevo.2018.00125.

Quinard A, and Cézilly F. 2012. Sex Roles during Conspecific Territorial Defence in the Zenaida Dove, Zenaida Aurita. Animal Behaviour 83 (1): 47–54. https://doi.org/10.1016/j.anbehav.2011.09.032.

Remage-Healey L, and Bass AH. 2004. Rapid, hierarchical modulation of vocal patterning by steroid hormones. Journal of Neuroscience 24 (26): 5892–5900. https://doi.org/10.1523/JNEUROSCI.1220-04.2004.

Remage-Healey L, and Bass AH. 2006. A rapid neuromodulatory role for steroid hormones in the control of reproductive behavior. Brain Research, Sex, Genes and Steroids, 1126 (1): 27–35. https://doi.org/10.1016/j.brainres.2006.06.049.

Ribble DO, and Salvioni M. 1990. Social organization and nest co-occupancy in *Peromyscus californicus* a monogamous rodent. Behavioral Ecology and Sociobiology 26: 9–16.

Rieger NS, and Marler CA. 2018. The function of ultrasonic vocalizations during territorial defence by pair-bonded male and female California mice. Animal Behaviour 135: 97–108. https://doi.org/10.1016/j.anbehav.2017.11.008.

Rieger NS, Stanton EH, and Marler CA. 2019. Division of labour in territorial defence and pup retrieval by pair-bonded California mice, *Peromyscus californicus*. Animal Behaviour 156: 67–78. https://doi.org/10.1016/j.anbehav.2019.05.023.

Rogers FD, Rhemtulla M, Ferrer E, and Bales KL. 2018. Longitudinal trajectories and inter-parental dynamics of prairie vole biparental care. Frontiers in Ecology and Evolution 6: 73. https://doi.org/10.3389/fevo.2018.00073.

Rogers W. 1988. Parental investment and division of labor in the midas cichlid (*Cichlasoma citrinellum*). Ethology 79 (2): 126–42. https://doi.org/10.1111/j.1439-0310.1988.tb00706.x.

Roozen HG, Boulogne JJ, van Tulder MW, van den Brink W, De Jong CAJ, and JFM Kerkhof. 2004. A systematic review of the effectiveness of the community reinforcement approach in alcohol, cocaine and opioid addiction. Drug and Alcohol Dependence 74 (1): 1–13. https://doi.org/10.1016/j.drugalcdep.2003.12.006.

Rosenfeld CS, Johnson SA, Ellersieck MR, and Roberts RM. 2013. Interactions between parents and parents and pups in the monogamous California mouse (*Peromyscus californicus*). PLOS ONE 8 (9). https://doi.org/10.1371/journal.pone.0075725.

Slabbekoorn H. 2013. Songs of the city: noise-dependent spectral plasticity in the acoustic phenotype of urban birds. Animal Behaviour, Including Special Section: Behavioural Plasticity and Evolution 85 (5): 1089–99. https://doi.org/10.1016/j.anbehav.2013.01.021.

Timonin ME, Kalcounis-Rueppell MC, and Marler CA. 2018. Testosterone pulses at the nest site modify ultrasonic vocalization types in a monogamous and territorial mouse. Ethology 124 (11): 804–15. https://doi.org/10.1111/eth.12812.

Tinbergen N. 1957. The functions of territory. Bird Study 4 (1): 14–27. https://doi.org/10.1080/00063655709475864.

Trainor BC, and Marler CA. 2001. Testosterone, paternal behavior, and aggression in the monogamous California mouse (*Peromyscus californicus*). Hormones and Behavior 40 (1): 32–42. https://doi.org/10.1006/hbeh.2001.1652.

Trainor BC, and Marler CA. 2002. Testosterone promotes paternal behaviour in a monogamous mammal via conversion to oestrogen. Proceedings of the Royal Society B: Biological Sciences 269 (1493): 823–29. https://doi.org/10.1098/rspb.2001.1954.

Trainor BC, Bird IM, and Marler CA. 2004. Opposing hormonal mechanisms of aggression revealed through short-lived testosterone manipulations and multiple winning experiences. Hormones and Behavior 45 (2): 115–21. https://doi.org/10.1016/j.yhbeh.2003.09.006.

Wingfield JC, Hegner RE, Dufty AM, and Ball GF. 1990. The ‘Challenge Hypothesis’: theoretical implications for patterns of testosterone secretion, mating systems, and breeding strategies. The American Naturalist 136 (6): 829–46. https://doi.org/10.1086/285134.

Wingfield JC, Jacobs J, Tramontin AD, Perfito N, Meddle SL, Maney DL & Soma KK. 1999. Toward an ecological basis of hormone-behavior interactions in reproduction in birds. in Reproduction in context: environmental and social influences on reproductive behavior and physiology. MIT Press, pp. 85–128.

Wingfield JC, Goymann W, Jalabert C, and Soma KK. 2020. “Reprint of ‘Concepts Derived from the Challenge Hypothesis.’” Hormones and Behavior 123: 104802. https://doi.org/10.1016/j.yhbeh.2020.104802.

Wood RI. 2004. Reinforcing aspects of androgens. Physiology and Behavior, Male Sexual Function, 83 (2): 279–89. https://doi.org/10.1016/j.physbeh.2004.08.012.

Zhang F, Chen P, Chen Z, and Zhao J 2015. Ultrasonic frogs call at a higher pitch in noisier ambiance. Current Zoology, 61, 996–1003. https://doi.org/10.1093/czoolo/61.6.996

Zhao X, and Marler CA. 2014. Pair bonding prevents reinforcing effects of testosterone in male California mice in an unfamiliar environment. Proceedings. Biological Sciences 281 (1788): 20140985. https://doi.org/10.1098/rspb.2014.0985.

Zhao X, and Marler CA. 2016. Social and physical environments as a source of individual variation in the rewarding effects of testosterone in male California mice (*Peromyscus californicus*)” Hormones and Behavior 85: 30–35. https://doi.org/10.1016/j.yhbeh.2016.07.007.

Zhao X, Fuxjager MJ, McLamore Q, and Marler CA. 2019. Rapid effects of testosterone on social decision-making in monogamous California mice (*Peromyscus californicus*). Hormones and Behavior 115, https://doi.org/10.1016/j.yhbeh.2019.06.008.

Zhao X, Castelli FR, Wang R, Auger AP, and Marler CA. 2020. Testosterone-related behavioral and neural mechanisms associated with location preferences: a model for territorial establishment. Hormones and Behavior 121: 104709. https://doi.org/10.1016/j.yhbeh.2020.104709.

Ziegler TE, Washabaugh KF, Snowdon CT. 2004. Responsiveness of expectant male cotton-top tamarins, *Saguinus oedipus,* to mate’s pregnancy. Hormones and Behavior;45:84–92. https://doi.org/10.1016/j.yhbeh.2003.09.003.

